# Coupling allows robust redox circadian rhythms despite heterogeneity and noise

**DOI:** 10.1101/2023.02.12.528191

**Authors:** Marta del Olmo, Anton Kalashnikov, Christoph Schmal, Achim Kramer, Hanspeter Herzel

## Abstract

Circadian clocks are endogenous oscillators present in almost all cells that drive daily rhythms in physiology and behavior. There are two mechanisms that have been proposed to explain how circadian rhythms are generated in mammalian cells: through a transcription-translation feedback loop (TTFL) and based on oxidation/reduction reactions, both of which are intrinsically stochastic and heterogeneous at the single cell level. In order to explore the emerging properties of stochastic and heterogeneous redox oscillators, we simplify a recently developed kinetic model of redox oscillations to an amplitude-phase oscillator with ‘twist’ (period-amplitude correlation) and subject to Gaussian noise. We show that noise and heterogeneity alone lead to fast desynchronization, and that coupling between noisy oscillators can establish robust and synchronized rhythms with amplitude expansions and tuning of the period due to twist. Coupling a network of redox oscillators to a simple model of the TTFL also contributes to synchronization, large amplitudes and fine-tuning of the period for appropriate interaction strengths. These results provide insights into how the circadian clock compensates randomness from intracellular sources and highlight the importance of noise, heterogeneity and coupling in the context of circadian oscillators.

## I. Introduction

Circadian clocks are endogenous oscillators that control daily rhythms in metabolic, physiological and behavioral processes in accordance with periodically reocurring environmental zeitgebers [1]. Although the molecular ‘core’ components have widely different evolutionary origins, a common design principle is present in almost all organisms where circadian timekeeping mechanisms have been investigated [2–5]. This scheme relies on a number of interlocked transcription-translation negative feedback loops (TTFLs) and additional regulatory mechanisms such as epigenetic [6] or post-translational modifications [7]. Since circadian clocks are cell-autonomous, even single cells oscillate with self-sustained circadian periods [8–11]. These intrinsic rhythms also synchronize to the outer world to maintain coherent physiological rhythms [1, 3].

In mammals, the ‘canonical’ cell-autonomous molecular timekeeping machinery [4, 12] consists of the positive regulators CLOCK and BMAL1 that induce the expression of a manifold of genes, including the negative regulators PERs and CRYs. These proteins form large macromolecular complexes [13] that translocate to the nucleus to repress their own expression by inhibiting the CLOCK:BMAL1 activity at their promoters. This process generates a cell-autonomous self-sustaining cycle of gene and protein expression that occurs throughout a 24 h period. But there is accumulating evidence from the last decades suggesting that transcription-independent processes can also generate circadian rhythmicity. These non-canonical oscillator mechanisms include the cyanobacterial KaiABC phosphorylation clock [14] or the circadian oxidation of peroxiredoxin (Prx) proteins [15], a ubiquitous family of antioxidant enzymes which are active in their reduced state and that contribute to removal of reactive oxygen species.

Oxidation of Prx proteins has been found to contribute to circadian timekeeping in human red blood cells, which are devoid of nuclei or ribosomes and therefore cannot oscillate according to the canonical TTFL [15]. Although the molecular mechanisms behind Prx oxidation rhythms differ across cell types and species, these rhythms seem to be conserved across all kingdoms of life [16], in contrast to the divergent evolution of the TTFL. In mouse red blood cells, oxidation of Prx occurs to ‘clean’ the erythrocyte from H_2_O_2_ generated as a result of hemoglobin auto-oxidation, and oxidized Prx is then degraded by the 20S proteasome [17]. In mitochondria from heart, adrenal gland and brown adipose tissue, however, the oxidized Prx is not degraded but reduced back to its active state by sulfiredoxin (Srx). In these tissues, Srx and hyperoxidized Prx3 undergo antiphasic circadian oscillations [18] and contribute to rhythms of H_2_O_2_.

We previously generated the first kinetic model that describes mitochondrial Prx3/Srx oscillations [19]. Nevertheless, it is widely recognized that a deterministic description of genetic and protein regulatory networks may be questionable, because protein molecules involved in these regulatory mechanisms act at rather low concentrations in single cells [20]. Cellular processes are intrinsically stochastic and very heterogeneous at the single-cell level [21–23] and therefore we hypothesized that the molecular processes happening in single mitochondria ought to be noisy too. For this reason, to explore generic properties of noisy and heterogeneous clocks, we simplified the kinetic model to a stochastic amplitude-phase redox oscillator. Additionally, we investigate how different redox oscillators interact with each other and to a TTFL to minimize the impact of noise and heterogeneity.

Here, we study ensembles of redox amplitude-phase oscillators with amplitude-period correlations (a phenomenon known as ‘twist’) subject to Gaussian noise. We show that noise and heterogeneity alone lead to desynchronization, consistent with the damping of rhythms observed in extracted tissues [24] or in cell cultures after a synchronizing stimulation [9, 10, 25]. We find that, when all oscillators are identical and run at the same speed, intrinsic oscillator properties influence the dynamics of noise-induced desynchronization: rigid oscillators are more robust, while more ‘plastic’ oscillators desynchronize faster. If oscillators differ in their free-running periods, this heterogeneity contributes to a faster desynchronization. We then consider the interaction of coupling among mitochondrial oscillators through a mean-field and coupling of the ensemble to a TTFL system of a slightly different period. We find that both mean-field coupling as well as coupling to a TTFL compensate the noise- and heterogeneity-induced desynchronization and can induce amplitude expansions as well as tune the periods of the individual oscillators, establishing synchronized rhythms for appropriate interaction strengths. The simulations in this work highlight the importance of heterogeneity and noise in population timing, demonstrate their complex interactions and illustrate how mean-field coupling or timing cues from an additional clock can help to promote synchrony among non-identical noisy oscillators. These findings contribute to the understanding of coupled oscillator networks, from single organelles to tissues and organisms, and how coupling contributes to robust rhythmicity.

## II. Results

### A stochastic phenomenological model of circadian redox oscillators

In our previous study, we introduced the first kinetic model of ordinary differential equations (ODEs) that describes circadian redox oscillations of hyperoxidized Prx3 (Prx3-SO_2_H) and Srx in mitochondria [19]. A detailed review of Prx3 structure, mechanism and function can be found in [18, 26]. In brief, mitochondrial H_2_O_2_ levels (what we call ‘danger 1’ or *D*_1_) are controlled by Prx3 in its active state *A* (Figure 1A). But *A*, however, can become catalytically inactive through *D*_1_-mediated hyperoxidation, resulting in the inactive Prx3 *I*. As *A* is converted to *I* and the Prx3 pool becomes inactive, the peroxide *D*_1_ accumulates in the mitochondrion and can overflow to the cytosol, where it is referred to as ‘danger 2’ or *D*_2_. In the cytoplasm, *D*_2_ activates pathways to control its own production and, among others, it stimulates Srx oxidation and its import to mitochondria. Srx (the ‘rescuer’ *R*) then can reduce *I* back to *A*, constituting a negative feedback loop and thus ‘rescuing’ *I* and allowing a new cycle of Prx3 inactivation and H_2_O_2_ accumulation to start. We identified this scheme (Figure 1A, colored variables) as the minimal backbone that the system requires in order to oscillate in a self-sustained manner [19] with the characteristics that have been observed experimentally: a circadian period [18], a phase difference between hyperoxidized Prx3 and Srx of 8–12 h [18] and kinetic constants of the oxidation reactions in line with the reactivity reported in *in vitro* biochemical assays [26–28].

**Figure 1:**
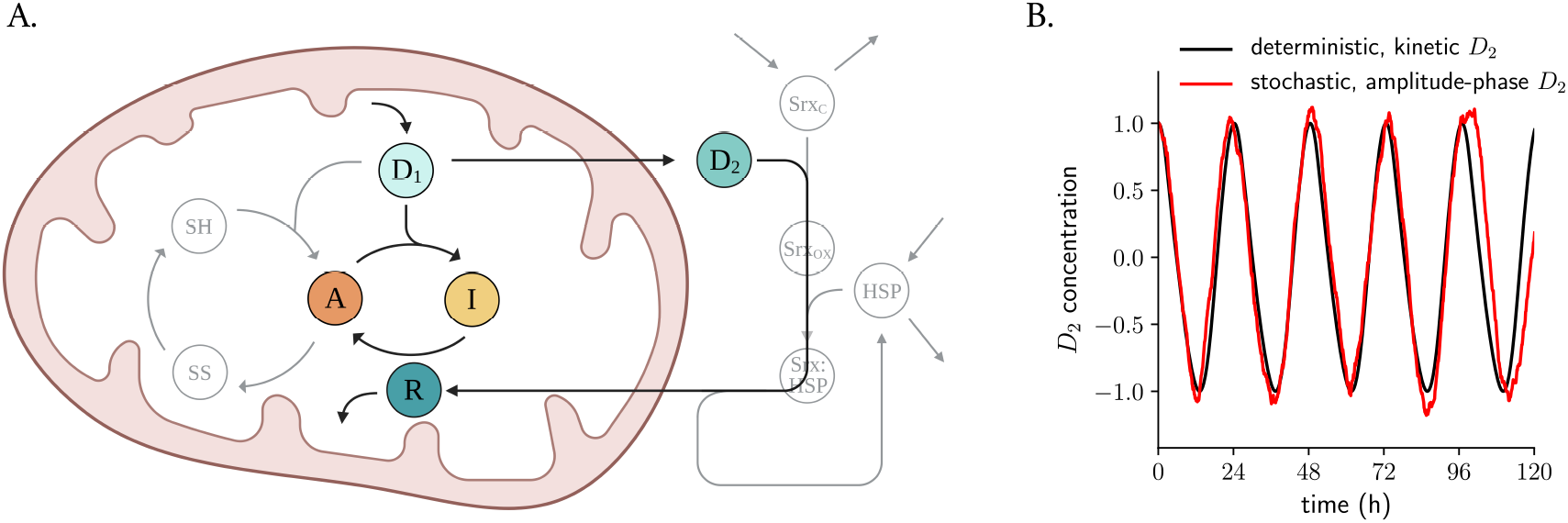
An amplitude-phase model with twist for stochastic circadian redox oscillations. **A**. Extended kinetic model and minimal backbone for circadian redox oscillations. Mitochondrial H_2_O_2_ levels (*D*_1_) are controlled by Prx3. The fully reduced state of the enzyme (-SH) eliminates *D*_1_ by reducing it to water and in turn oxidizing itself to an intermediate (-SOH) that we refer to as *A*. This oxidation is followed by a conformational change in which the -SOH group condenses with another Prx3 molecule, forming a disulfide bond (-SS); and finally, the homodimer can be reduced back allowing a new cycle to begin. A small fraction of Prx3, however, can become catalytically inactive through *D*_1_-mediated hyperoxidation of the -SOH intermediate (*A*) to Prx3-SO_2_H (*I*). As the hyperoxidation and inactivation reaction occurs, *D*_1_ accumulates in the mitochondrion and overflows to the cytosol. In the cytoplasm, H_2_O_2_ (now *D*_2_) stimulates Srx oxidation and complex formation with a heat shock protein Hsp90 that shuttles Srx to mitochondria, where it acts as the ‘rescuer’ *R* and reduces *I* back to *A*. The colored variables represent the minimal backbone that the system needs to oscillate with the experimentally observed characteristics, but the full model (including grey variables) oscillates along the same lines. **B**. *D*_2_ time series from the kinetic deterministic ODE-based model from (A) (black line) and from the stochastic simplified amplitude-phase oscillator (red line, see parameter estimation in Materials and Methods).

Modeling cell physiology with ordinary differential equations presumes a ‘deterministic’ view of the molecular interactions and translocation processes within cells. However, taking into account the stochastic fluctuations in the abundance of molecules and movements within mitochondria, it seems likely that these organelles behave as noisy (rather than deterministic) oscillators. For this reason, to study generic properties of noisy individual redox oscillators, we converted the kinetic deterministic model into a stochastic Poincaré oscillator model with twist (equation 1 in Materials and Methods). Poincaré oscillator models are simple descriptions of stable limit cycles in a 2-dimensional plane, and are one of the most abstract yet intuitive class of models, with only two variables, amplitude and phase, that have been widely used in chronobiology [29–36]. The equations do not depend on molecular details and can describe phenomenologically the oscillatory dynamics of a system [29, 37].

We simulated a stochastic amplitude-phase oscillator with twist and cycling at a 24.2 h period as in the kinetic model from [19] using the Euler-Maruyama method [38, 39]. The twist parameter *ϵ* was included to model the interdependence between oscillator period and amplitude, common in nonlinear oscillators [23, 40, 41], that we found in *D*_2_ oscillations: those with shorter periods are associated with smaller amplitudes (Materials and Methods, Supplementary Figure S1A). The amplitude relaxation rate parameter *λ* was calculated from simulations in which we applied perturbations at random phases and then computed the rate at which the perturbations relax back to the limit cycle (Materials and Methods, Supplementary Figure S1B).

With this parametrization we converted the kinetic model to a Poincaré oscillator model specific to the redox system and added different noise intensities to the system. As expected, we found that noise contributes to damping of the average signal of 100 oscillators (Supplementary Figure S2A), with higher noise intensities desynchronizing the network faster (Supplementary Figure S2B). Noise intensity was fixed for the rest of the simulations at a standard deviation *σ*_*x*_ = 0.05. At the level of individual oscillators, however, we found that our Poincaré model captures the oscillatory properties of the deterministic *D*_2_ dynamics despite the white noise component (Figure 1B): analysis in the power spectrum revealed a peak at 24.2 h (Supplementary Figure S2C), as expected from the period of the deterministic *D*_2_ oscillations. The autocorrelation function of an individual noisy oscillator (the correlation of one noisy amplitude-phase oscillator with itself, but after a time delay) decays exponentially over time due to the noise component (Supplementary Figure S2D).

### Noise and oscillator heterogeneity desynchronize a network of redox oscillators

Mitochondria vary in number depending on the cell type: while a human liver cell contains 1 000–2 000 mitochondria, a cardiac myocyte can have up to 10 000 [42]. To analyze the effect of noise on an ensemble of mitochondria, we simulated 100 identical stochastic *D*_2_ noisy amplitude-phase oscillators. Despite all oscillators starting with the same initial conditions (and thus same phase), they drift apart in phase over time due to the white noise component (Figure 2A, B, Supplementary Figure S2A). We computed the average standard deviation of the phase dispersion over time across different simulations and observed, surprisingly, that the phase spreading does not grow with the square-root of time (Figure 2C, red curve) as expected from Fick’s Second Law of Diffusion and other theoretical studies [39, 43, 44]. When exploring the parameter space of the stochastic model, we found that higher values of amplitude relaxation rate *λ* result in the pure phase diffusion, i.e. in the standard deviation of phase dispersion growing over time according to Fick’s predicted square-root function (Supplementary Figure S3A). These results provide an intuitive explanation to why *λ* influences the desynchronization dynamics: noisy oscillators, if rigid (high *λ* values), undergo less amplitude excursions because of the high relaxation rate that ‘pulls’ them back to the *A* = 1 default state. But in weaker oscillators with lower values of *λ*, however, the inherent noise component contributes to larger amplitude fluctuations. Moreover, the explicit twist *ϵ* of our model system makes from any amplitude fluctuation an inevitable phase shift, what results in faster and more complex desynchronization dynamics for weak oscillators that no longer fit to a square root.

**Figure 2:**
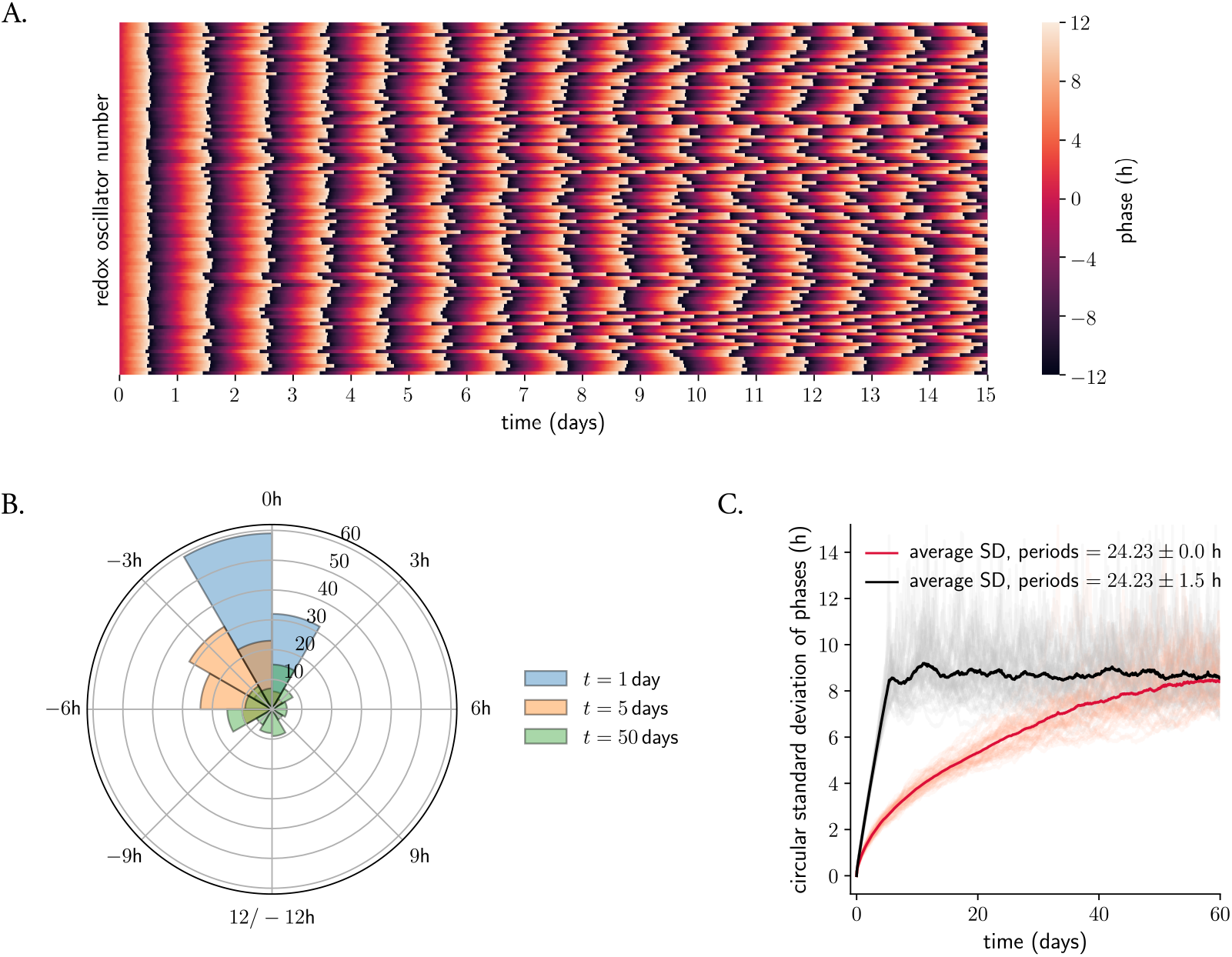
Noise and heterogeneity contribute to desynchronization and phase dispersion of redox oscillators. **A**. Desynchronization of 100 identical amplitude-phase redox oscillators over time. Colors indicates the instantaneous phase *ϕ*_*i*_ of each oscillator (determined as *ϕ*_*i*_ = arctan 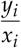, see Materials and Methods), normalized to a 24 h period. **B**. Phase dispersion over time in a network of 100 identical amplitude-phase stochastic redox oscillators at 1, 5 and 50 days. **C**. Period heterogeneity enhances phase dispersion: different desynchronization dynamics for an ensemble of identical (red) or heterogeneous (black) amplitude-phase stochastic redox clocks. The thick red/black lines indicate the average standard deviation from 50 different realizations of network simulations; each realization (light red/grey) was composed of a network of 100 oscillators, as in (A). Phases are normalized to a 24 h cycle.

An assumption that is commonly made is that homogeneous systems and those with slight heterogeneity will behave similarly. We challenged this hypothesis by introducing heterogeneity in each oscillator’s period and analyzing the desynchronization dynamics of the ensemble. Period values were taken from a normal distribution with mean = 24.23 h and standard deviation = 1.5 h (studies in dispersed –and presumably uncoupled– SCN neurons or U2-OS cells have estimated standard deviations of 1.5–2 h [8, 25, 45]). Redox clocks desynchronize faster when they are modeled as ensembles of noisy and heterogeneous oscillators compared to noisy identical oscillators: networks of heterogeneous clocks (Figure 2C, black curve) switch faster to high standard deviation values of phase dispersion as compared to identical oscillators (red curve). We had previously seen that rigid oscillators (those with high values of *λ*) are more resistant to noise-induced desynchronization (Supplementary Figure S3A). This dependence was weakened when heterogeneity was introduced in the ensemble: heterogeneity in oscillator period at a standard deviation = 1.5 h controls the speed of desynchronization of the ensemble, independent of amplitude relaxation rate values (Supplementary Figure S3B).

### Coupling counteracts noise- and heterogeneity-induced desynchronization

Networks of coupled oscillators are widespread throughout the living and the non-living worlds [46, 47]. Two or more oscillators are said to be coupled if some physical or chemical process allows them to influence one another. We next speculated that, if single mitochondria are competent noisy redox oscillators, they likely exchange time information that leads to the adjustment of individual phases (and periods) towards a common phase (and period) in the cellular environment where they tick. Clear bands of Prx-SO_2_ in western blots from red blood cells support this assumption: although these cells have no mitochondria, if their Prx redox cycles were not coordinated, oscillators would dephase over time (as in Figure 2) and it would not be possible to capture a robust average signal *in vitro*. We hypothesized that, in our model applicable to heart, adrenal gland and brown adipose tissue, coupling between mitochondria may depend on the concentration of the synchronizing factor H_2_O_2_ (*D*_2_) in a way that, all organelles, through Srx (*R*), sense an ‘effective’ average concentration of all H_2_O_2_ molecules. The release of *D*_2_ (or *x*_*i*_ in the stochastic model) is expected to occur quickly relative to the 24 h oscillation cycle [48, 49]. As a result, *D*_2_ becomes evenly distributed in the cytoplasm, and all *x*_*i*_ oscillations can be approximated with an average level of all *x*_*i*_ signals or mean-field *M*. This mean-field then directly affects each individual oscillator *x*_*i*_ at a specific coupling strength *K*_*c*_ (equations 2 and 3, see Materials and Methods for details).

To understand the impact of increasing coupling strength on an ensemble of noisy redox oscillators, we analyzed the dynamics of a coupled ensemble (equations 3 in Materials and Methods) for three representative values of coupling strengths. In these simulations, all oscillators were assumed to have the same noise intensity *σ*_*x*_ = *σ*_*y*_ = 0.05, amplitude *A* = 1, amplitude relaxation rate *λ* = 0.05 h^-1^ and twist *ϵ* = 0.05 h^-1^ (see Materials and Methods), but to be heterogeneous in their periods, which were taken from a Gaussian distribution at 24.23 *±* 1.5 h. For no coupling (Figure 3A, top panel), no order is observed: all oscillators run at (or close to) their own intrinsic frequency –a state known as an ‘incoherent state’ in oscillator theory [29,50]. If the coupling strength is increased, order emerges: a fraction of the ensemble starts to run with a shared frequency, resulting in a substantial increase of the mean-field oscillation amplitude (black line in Figure 3A, middle panel). In this mixed state, partial synchronization occurs: a cluster of synchronized oscillators coexists with a fraction of non-synchronized oscillators, and the number of clocks that fall in the synchronized cluster increases with *K*_*c*_ (Figure 3A, bottom panel), consistent with previous theoretical studies [29]. Moreover, it is evident from the time series in Figure 3A how the amplitude of individual oscillators is also modulated by increasing coupling strengths, a phenomenon, which in the words of oscillator theory is known as ‘amplitude resonance effect’. In summary, coupling can establish synchronized rhythms and induce amplitude expansions of individual oscillators, directly affecting the mean-field signal.

**Figure 3:**
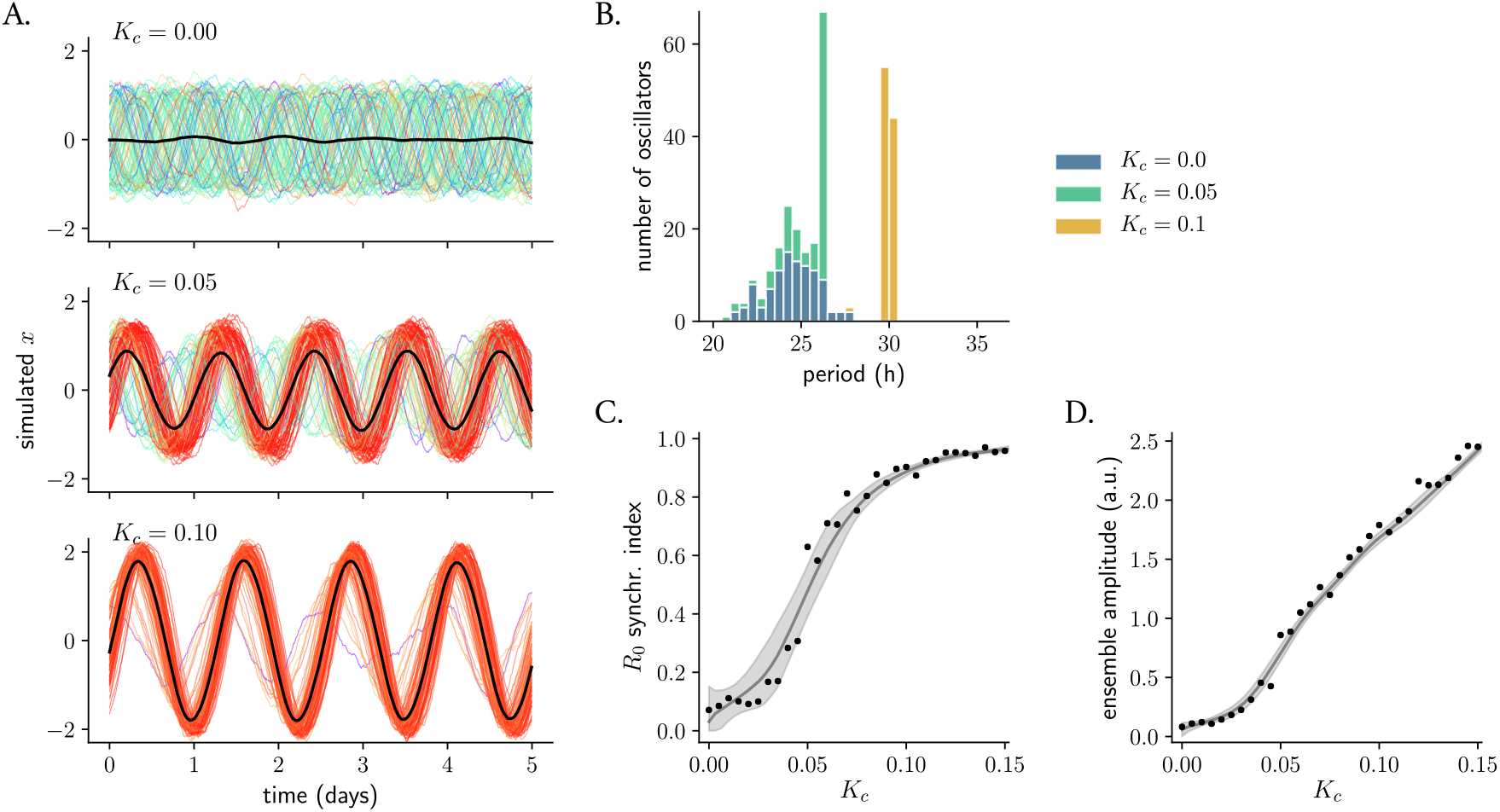
Coupling counteracts desynchronization by noise and period heterogeneity. **A**. Formation of temporal order upon coupling. Shown are numerical solutions of an ensemble of 100 stochastic and heterogeneous redox oscillators for three representative values of coupling strength. Numerical solutions of individual oscillators are color-coded with respect to their intrinsic period; the mean-field signal is highlighted in black. **B**. Histograms of periods of the individual oscillators from (A). Periods were determined by computing the zeroes of the oscillations and calculating the distance between two consecutive zeros with a negative slope. **C**. Network phase coherence, quantified with the order parameter *R*_0_ (equation 4 in Materials and Methods) as a function of the coupling strength *K*_*c*_. Instantaneous phases *ϕ*_*i*_ and amplitudes *A*_*i*_, used to compute *R*_0_, were determined as *ϕ*_*i*_ = arctan 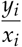 and by 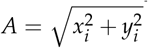 (with results validated with Hilbert transformation, data not shown). **D**. Amplitude of the ensemble in dependence of *K*_*c*_. Amplitude of the mean-field was determined as the half of the peak-to-trough distance, averaged across the last 8 oscillations after 100 days of simulation. In (C) and (D): points indicate the mean *R*_0_ or mean ensemble amplitude of 5 different realizations of network simulations; the solid line represents the average of 100 Lowess models that were fitted to 50% of the points (chosen randomly in each iteration); the grey shaded areas indicate the confidence interval for the Lowess models.

We next studied in more detail the rhythmic properties of the individual oscillators and the ensemble, namely periods, phases and amplitudes, for the three representative values of mutual coupling. For no coupling strength, the periods of the individual oscillators fall in a normal distribution with a mean period of 24.5 h and a standard deviation of 1.4 h (Figure 3B, blue bars), consistent with the known input period distribution of 24.23 *±* 1.5 h. But when a certain critical coupling strength is reached, a fraction of oscillators whose intrinsic period is close to the period of the mean-field starts to form a period-locked cluster, a phenomenon that has been previously referred to as ‘frequency pulling’ [51]. As predicted from previous computational studies [29], the number of oscillators in the period-locked cluster increases with higher coupling strengths (Figure 3B) and, as a consequence, the variance of the period distribution decreases. Note the tendency of redox oscillators to run at slower paces (larger periods) as the coupling strength is increased (Figure 3B): this is due to the intrinsic period-amplitude correlation (twist) with which the system is modeled, which makes from an amplitude expansion an inevitable period change (this will be discussed further in the next section and in the Discussion).

Increasing coupling strengths induces a re-organization of the oscillation amplitudes and phases that we quantified with the order parameter or synchronization index *R*_0_ [29, 50] (equation 4 in Materials and Methods). Instantaneous phases of the noisy and heterogeneous redox oscillators are equally distributed among their whole domain for no coupling strength (Figure 2B), but the synchronization index grows with increasing *K*_*c*_ (Figure 3C) resulting in phases clustered together in the oscillating ensemble. Note though that, sometimes, despite full synchronization with respect to periods, the phases and amplitudes of individual oscillators differ, what leads to an *R*_0_ *<* 1 (this is what happens for the particular case of *K*_*c*_ = 0.10, Figure 3A bottom panel and Figure 3C). On a population level, these different phases are termed chronotypes and they maintain a fixed angle with respect to the zeitgeber known as phase angle of entrainment. However, when comparing phases to the mean-field signal, these cannot be strictly called chronotypes, but rather ‘earlier’ and ‘later’ redox oscillators. When the ensemble is modeled without individual period differences, phase synchrony is achieved for lower values of *K*_*c*_ (Supplementary Figure S4). Furthermore, the amplitude of the mean-field is also expanded with *K*_*c*_ (Figure 3D) due to resonance effects. Because of the system’s positive twist and as previously mentioned, the period of redox oscillators also increases.

In summary, coupling can establish synchronized rhythms by coordinating phases, periods and amplitudes of individual oscillators, thus directly affecting the robustness of the mean-field signal. Amplitude modulations of individual oscillators due to resonance have an effect on the free-running periods, ultimately contributing to the period lengthening of the mean-field due to twist effects.

### Intrinsic individual oscillator properties affect the coupled network

Resonance effects are not only induced with increasing coupling strengths *K*_*c*_, but they also depend on intrinsic properties of the individual oscillators. First, the critical coupling strength needed to achieve synchronization increases with oscillator period heterogeneity (Supplementary Figure S4), similar to what has been observed in a Kuramoto model [29]. Furthermore, we observed that increasing amplitude relaxation rates *λ* has considerable effects in the amplitude expansion of the mean-field due to resonance. Rigid oscillators (e.g. with *λ* = 1 h^-1^, which return very fast to the stable limit cycle after a perturbation) show little variations in their amplitude as coupling strength increases, whereas more ‘plastic’ oscillators (e.g. with our default *λ* = 0.05 h^-1^) exhibit larger amplitude expansions (compare Figure 3A and Supplementary Figure S5A, also Supplementary Figure S5B). This is consistent with previous theoretical and experimental studies of coupling [29] and entrainment [52]. These studies have used the term ‘weak’ clocks for those with low amplitude relaxation rates *λ* but we rather call them ‘plastic’ oscillators indicating that their rhythmic properties can easily be shaped or modulated.

Together with rigid oscillators exhibiting lower amplitude expansions, we observed that they are more resistant to coupling-induced period differences. Whereas the period of the mean-field of a coupled and plastic ensemble shifted to values of 28 to 32 h for a representative value of *K*_*c*_ = 0.10, a rigid ensemble with *λ* = 1 h^-1^ stayed running at a pace of 24.8 h (Supplementary Figure S5C). Taken together, these results suggest that oscillators with higher amplitude relaxation rates are more robust to twist-induced effects. This is because the large relaxation rate value does not allow for large amplitude expansions upon coupling and, as a result, the period also remains stable.

### TTFL input further synchronizes an ensemble of weakly-coupled redox oscillators

Up until now in this study (and also in [19]), we have assumed that the production of H_2_O_2_ is constant. However, the sources of mitochondrial H_2_O_2_ in tissues where the redox clockwork is present have been shown to be partly under the control of the circadian TTFL. For example, in heart or brown adipose tissue, where oxidative metabolism is very high, the respiratory component represents a significant source of H_2_O_2_, and the rate of respiration has been shown to be rhythmically controlled by the canonical TTFL [54]. In adrenal gland, the main source of H_2_O_2_ is steroidogenesis (synthesis of corticosterone from cholesterol), which is regulated both by circadian pituitary release of ACTH [55–57] and by the adrenal clock [58]. Moreover, earlier studies have even shown that the rate-limiting enzyme in cholesterol production, StAR, has an E-box in its promoter [59] which could be potentially activated by a CLOCK:BMAL1 complex.

To explore the effects of a weakly-coupled redox oscillating system being influenced by the TTFL, we modified our redox ensemble to include a CLOCK:BMAL1 periodic signal from a Goodwin-like model [53], that we assumed to contribute to the production of H_2_O_2_ (Figure 4A, equations 5, 6 in Materials and Methods). Of note, the Goodwin-like model oscillated with a 23.6 h period (reproducing the results from [53]). Our simulations show that, when the intensity of the TTFL is strong enough, it can impose its 23.6 h period on the ensemble of weakly-coupled redox oscillators, effectively entraining them. Decomposition of the mean-field signal in its Fourier components nicely illustrates how the frequency of the TTFL (23.6 h) gains more weight in the redox mean-field signal as the TTFL input strength is increased (Figure 4B, left panel). In the absence of a TTFL, the weakly-coupled redox ensemble oscillates at a 26.2 h period (Figure 3A and Figure 4B, left and middle panels –orange line), but a TTFL input = 0.25 is able to entrain the redox ensemble, making it run at a 23.6 h period (Figure 4B, left and middle panels –green line). For intermediate TTFL inputs (between ∼ 0.05 and 0.09), however, not all of the redox oscillators entrain to the TTFL and the result is a more complex dynamics of the mean-field signal (and individual oscillators) and the observation of a phenomenon commonly referred to as ‘beating’ or ‘period-splitting’ in the time series (Figure 4B, right panel). Here, the frequency of the weakly-coupled mean-field co-exists and interacts with the frequency of the TTFL. These results highlight the importance of a proper interaction strength for successful entrainment to generate a robust signal in the network of weakly-coupled redox clocks.

**Figure 4:**
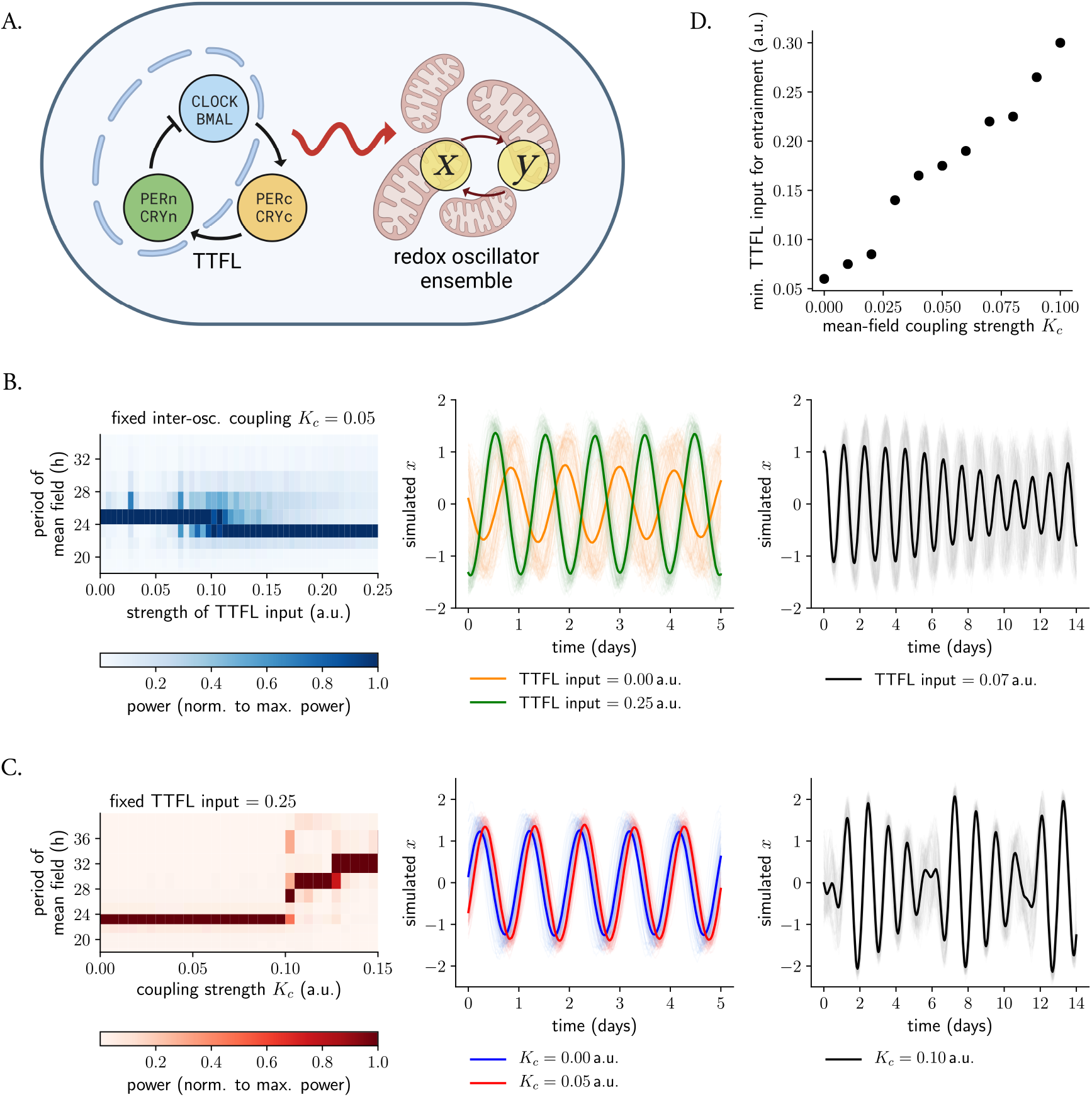
A TTFL input can further synchronize and entrain an ensemble of weakly-coupled redox oscillators. **A**. Scheme of the coupled TTFL-redox system. A Goodwin-like model [53], oscillating at a 23.6 h period, was used to drive an ensemble of 100 weakly-coupled stochastic amplitude-phase redox oscillators. **B**. A regulated TTFL input can entrain networks of weakly-coupled redox oscillators. Variations in period of the redox mean-field as a function of the strength of the TTFL input for a fixed value of coupling strength *K*_*c*_ (left panel): when the TTFL input is strong enough, it can impose its 23.6 h period in the network of weakly-coupled oscillators (middle panel, green line), but in the absence of the TTFL, the ensemble oscillates at the mean-field 26.2 h period (middle panel, orange line). For intermediate TTFL input strengths, no entrainment occurs and beating signals due to co-existing frequencies are observed (right panel). **C**. Increasing coupling strengths hinder TTFL entrainment. Variations in period of the redox mean-field as a function of the strength of the inter-oscillator coupling strength *K*_*c*_ for a fixed TTFL input (left panel): the higher the coupling, the more robust the network is to TTFL entrainment (and instead oscillates at the frequency of the mean-field). Representative entrained timeseries are shown for a fixed TTFL input and two mean-field coupling strengths (middle panel, *K*_*c*_ = 0.00 in blue; *K*_*c*_ = 0.05 in red). Complex dynamics occur due to co-existing frequencies for high coupling strength *K*_*c*_, which make the system more robust to TTFL entrainment (right panel). In the left panels of (B) and (C), periods were calculated from periodograms of the mean-field; colors indicate the normalized power of each frequency. In the middle and right panels of (B) and (C), individual oscillators are shown in light; the mean-field signal is shown with thicker lines. **D**. Minimum TTFL input neeeded for entrainment as a function of the coupling strength *K*_*c*_: networks of redox oscillators with higher *K*_*c*_ values need stronger TTFL inputs to entrain.

Mitochondrial redox clockworks might be coupled to different extents in different tissues. To explore how changing mean-field coupling strengths affect the entrainment of the redox network to a TTFL, we simulated networks with different *K*_*c*_ values all being driven by a TTFL at an input = 0.25 (which we previously found to entrain a weakly-coupled ensemble, Figure 4B). We observed that the TTFL only entrains redox ensembles when their inter-oscillator coupling is under a threshold value: strongly coupled networks (with *K*_*c*_ *>* 0.11 in Figure 4C, left panel) do not entrain to the TTFL, suggesting that coupling makes these oscillators more rigid.

In line with this observation, we found that the minimum TTFL input needed for entrainment increases with coupling strength *K*_*c*_ (Figure 4D). Mean-field amplitudes of networks of weakly-coupled redox oscillators entrained to a TTFL input (Figure 4C, middle panel, red line) were comparable to those from networks of uncoupled redox oscillators driven by the canonical TTFL (blue line). Moreover, increasing *K*_*c*_ also led to the observation of complex dynamics due to interacting frequencies (Figure 4C, right panel). These results suggest that, while coupling might not contribute majorly to more robust rhythmicity and higher amplitudes in ensembles of already entrained redox oscillators, it does give the system ‘timing autonomy’ in case that the TTFL fails or is somehow disrupted.

## III. Discussion

The presence of noise in enzymatic and genetic networks is a fundamental consequence of the stochastic nature of biochemical reactions due to the low numbers of molecules and the local environmental fluctuations that are present in single cells. The ability to function effectively and consistently ensuring reliable transmission of information amidst such random fluctuations is a major issue in network behavior and gene expression. In this paper we present a simple model leveraging the amplitude-phase equations with Gaussian noise to describe circadian redox oscillations at the level of single mitochondria. We show (i) that noise desynchronizes a network of redox oscillators and (ii) that differences between individual oscillators enhance desynchrony, but (iii) that coupling among redox oscillators and to a genetic network of a TTFL, however, can compensate for and filter the effects of biochemical noise and heterogeneity to create a synchronized, robust and high amplitude circadian redox system.

Even though the model from Figure 1A [19] describes the observed rhythmic properties of Prx and Srx oscillations [18], the system’s kinetic details remain unknown. Previous biochemical assays have estimated some of the relevant kinetic constants [26], but whether the same conditions are maintained in the cellular environment is not clear, and thus enzymatic affinities or kinetic constants might differ. To address this uncertainty, simplifying kinetic models to amplitude-phase models represents an advantage, as these models have fewer parameters and do not rely on molecular details. From all the variables from the kinetic model we focused on cytoplasmatic H_2_O_2_ (*D*_2_) oscillations because the endogenous levels of this molecule have been measured in a high-resolution manner with fluorescent reporters and because H_2_O_2_ has been shown to act as a signaling molecule [60] and hence is a candidate that might transmit timing information to the TTFL.

Converting the kinetic deterministic model into a stochastic amplitude-phase model allowed us to quantitatively analyze the emergent properties that arise from a network of stochastic clocks with different intrinsic oscillator properties in response to coupling to a mean-field or to a TTFL input. Noise in the Cartesian coordinates of the model was directly translated into the phases of individual oscillators dispersing, with the dynamics of phase dispersion depending on intrinsic oscillator properties. Pure phase diffusion, i.e. the standard deviation of phase dispersion growing with the square root of time, was only observed for rigid oscillators with high relaxation rates (Figure 2C, Supplementary Figure S3).

A phenomenon that has been underlooked in previous theoretical or experimental studies of circadian rhythms is that of twist. This term refers to amplitude-period correlations, and such inter-dependencies are generic features of nonlinear oscillators, from planets moving around stars [61] to molecular clocks of U2-OS cells [23] or of cells in the choroid plexus [41]. Since our self-sustained kinetic redox oscillator model is based on nonlinearities, co-variations of amplitudes and periods in the rhythms were expected. We indeed found a positive amplitude-period correlation (Supplementary Figure S1A) and hence we included an explicit twist parameter in our framework. We observed that twist is related to the robustness of the rhythmic ensemble: oscillators with higher amplitude relaxation rates are more resistant to coupling-induced twist effects and thus to period-lengthening or amplitude expansions due to resonance (Supplementary Figure S5B, C). Practically, this means that rigid limit cycles could be approximated with a phase oscillator, neglecting amplitude effects [40]. In our particular case of redox clocks with positive twist and low relaxation rate, the individual periods shift to larger values due to the resonance effects that come with increasing coupling. This implies that any observation of resonant behavior also provides information on the underlying oscillator type and on the presence of twist. For example, the systematic period increase and amplitude expansions that were observed in a theoretical study of synchronization of neurons in the SCN [53] can be justified with an implicit positive twist and with low oscillator relaxation rates.

It is important to remark that, throughout our work, we have used the term ‘rigid’ oscillators to describe those clocks whose rhythmic properties cannot be easily changed or adapted in responds to changes that the system encounters (e.g. in the face of mutual coupling or zeitgeber input). Previous theoretical studies, however, have quantified oscillator rigidity using Floquet theory [52, 62]: the higher the Floquet multiplier, the more rigid a system is. Both amplitude relaxation rate and coupling strength have been shown to contribute to larger Floquet multipliers [52, 62] and are thus intimately related to the rigidity of the oscillatory system (coupled or uncoupled) as a whole.

The role of coupling at the molecular level and its effects on the average signal are well established from both theoretical [29, 53] and experimental studies [25]. But, is a strongly coupled system more or less robust against perturbations? Rationale (1): on the one hand, it makes sense to reason that a strongly coupled system is very robust against perturbations. The more rigid the network is (i.e. the larger the coupling), the more difficult it is to perturb it and, hence, the smaller the phase shift caused by a zeitgeber will be. Thus, decreasing the inter-oscillator coupling would make the system more susceptible to perturbations (phase shifts) according to this hypothesis. Rationale (2): on the other hand, one might think that a weakly-coupled ensemble shows smaller responses to perturbations than a strongly coupled system. This can be argued from the view that, in a weakly-coupled network, individual oscillators are not fully synchronized and thus do not share the same phase at the time of perturbation. As a result, the response of the mean-field (the average response of all the oscillators) might be masked by the average of the individual responses. As the coupling between the oscillators increases, however, their responses to the perturbation become more aligned, resulting in a larger response of the mean-field to a perturbation.

Interestingly, there is experimental evidence suggesting that both contradicting hypotheses might hold. On the one hand and in support of ad (1), upon inhibition of TGF-*β* signaling (suggested to promote inter-cellular coupling in peripheral clocks) a temperature shock could shift the phase of circadian rhythms by about 8 h, in comparison with solvent-treated cells, where no significant phase shift was observed [25]. In addition, mice with impaired AVP signaling (a potent coupling factor in the SCN) recover faster from jet lag than wild-type mice [63]. In support of ad (2), however, there is data showing that highly synchronized SCN networks with high-amplitude rhythms show a larger phase-shifting capacity than desynchronized networks of low amplitude [64, 65]. More theoretical studies are needed to resolve these conflicting hypotheses and address differences, but it is possible that the relative importance of one effect over the other depends on the system being studied and the perturbation being applied. Our results, while they do not answer the question to robustness to perturbations applied at a certain time, predict that the higher the inter-oscillator coupling in a system, the more robust it becomes to entrainment (Figure 4D). In line with our findings, previous experimental studies on entrainment [52] have actually shown that lung clocks entrain to extreme zeitgeber cycles whereas SCN clocks (believed to have a higher inter-oscillator coupling) do not.

An important goal is to understand the relationship between redox clocks and the TTFL, as it is becoming evident that there is an interplay between both. Although it is clear that the TTFL regulates a manifold of metabolic and redox processes (see [66] for a nice review on the interplay between the clock and metabolism), redox oscillators seem ‘independent’ clocks that tick in an autonomous manner, although with altered properties when certain clock components are disturbed or absent. In behaviorally arrhythmic *Drosophila* mutants and *Neurospora crassa* mutants with lengthened periods, Prx oscillations are perturbed in phase [16]. Embryonic fibroblasts derived from *Cry1*^*-/-*^*;Cry2*^*-/-*^ mice show robust Prx3 rhythms but with increased period length [15]. Additionally, the TTFL has been shown to control a number of oxidative processes in adrenal gland [55–57] or heart [54], which might also explain why ensembles of mitochondria oscillate robustly *in vitro* and *in vivo* [18, 67].

To address this interplay, we coupled our network of self-sustained stochastic redox clocks to a TTFL, simulating a CLOCK:BMAL1 regulation of H_2_O_2_ generation. We reproduced the findings of [15], namely that the redox ensemble coupled to a TTFL oscillates with a ∼24 h period (entraining to the TTFL), but in the absence of the TTFL, the weakly-coupled network oscillates at a slightly lengthened 26.2 h period (Figure 4B). Moreover, our results suggest that whereas weak mean-field coupling does not have a major effect in a redox ensemble that is already entrained to a TTFL in terms of amplitude (Figure 4C), it does confer the redox ensemble certain autonomy to oscillate coherently and maintain its time-telling properties should the TTFL fail.

Not only is there evidence for the TTFL regulating redox clocks, but the H_2_O_2_ release might also represent a potential coupling node between both oscillators. Our kinetic Prx3/Srx redox oscillator model contains only one negative feedback, namely the reduction of *I* back to *A* by *R*, which is stimulated by the cytosolic H_2_O_2_ increase. Although cytosolic H_2_O_2_ has traditionally been regarded as a dangerous oxidant whose levels had to be be tightly controlled, the current paradigm is that it can signal as a second messenger and, for instance, activate the p38 MAPK pathway to decrease mitochondrial H_2_O_2_ production [18, 55]. The periodic H_2_O_2_ release to the cytosol is expected to act on other cytosolic or nuclear targets that have not been identified yet. Canonical clock proteins might be directly or indirectly affected by H_2_O_2_. This idea is supported by a study from 2014, that showed that the interaction of PER2 and CRY1 is redox-sensitive [68], as well as by other studies that have shown that the cellular redox poise can regulate the TTFL oscillator through NAD^+^ levels [69] or heme [70]. Moreover, H_2_O_2_ has been shown to oscillate robustly in cultured cells and mouse liver [67], and CLOCK contains a redox-sensitive cysteine that responds to changes in intracellular H_2_O_2_ [67]. These findings provide compelling evidence for a role of this oxidant in regulating the TTFL. The reciprocal interplay between the Prx system and the local TTFL clock might allow synchronization between local metabolic activity and systemic circadian regulation.

It is important to emphasize that organisms have evolved networks to function in extremely noisy cellular environments [20, 71, 72]. Coupling is a strategy that can confer resistance against such noise, but additional mechanisms might include switch-like events in spatial or temporal domains (bistability [73], ultrasensitivity [74]) or feedback regulation and suitable network designs [75–77], among others. In addition, some of these networks may not only be resistant to, but may also actively exploit, cellular noise to perform their functions under conditions in which it would not be possible by deterministic means. In certain cases, noise can act as a ‘driving force’ to, for instance, maintain or induce oscillations in a pathway. For example, in the case of weakly damped clocks, if the noise component is too low, the rhythms may become damped out over time and transmission of information might get compromised. However, if the noise is just right, it can help to ‘kick start’ the oscillations and maintain robust rhythms [78, 79]. There is actually an ongoing discussion whether circadian clocks in single cells are self-sustained limit cycles or weakly damped oscillators. If the latter, noise might certainly contribute to the generation of robust rhythms.

In general, the role of noise in oscillatory pathways is complex and depends on the specific details of the system. In our design, we have shown how it contributes to network desynchronization and how coupling can overcome the noise- and heterogeneity-induced desynchronization. Coupling is a characteristic property of the circadian clock timing system seen at all levels of organization. At the molecular level, interlocked feedback loops of clock proteins and genes are stabilized by post-translational mechanisms involving signaling components. At the cellular level, coupling contributes to robustness of tissue pacemakers by exchange of timing molecules. Ultimately, all tissues are coupled to generate daily physiological rhythms in response to, both, internal and external demands. Quantifying the noise present at the level of biochemical reactions as well as behavioral responses, and how it is buffered (or used) by a biochemical or genetic network might be helpful in order to understand how tissues maintain coherent rhythms in an organism. Overall, the complex interplay of coupling at the molecular, and inter-cellular levels, as well as coupling to physiological and environmental factors is likely what drives the robust and rhythmic behavior of circadian rhythms at the cellular, tissue and organismic levels.

## IV. Limitations of the Model and Conclusions

Building effective stochastic models requires a solid foundation of the preliminary deterministic model and substantial amounts of additional quantitative data about cell constituents and cell behavior (reviewed in detail by [71, 72, 80]) to reliably estimate noise properties. Unfortunately, quantitative data at the level of mitochondrial H_2_O_2_, Prx3 and Srx rhythms is still scarce, and hence noise was introduced heuristically by adding a Wiener process to the *x* and *y* coordinates of the redox amplitude-phase oscillator. Period-amplitude correlations have been experimentally observed in rhythms of the TTFL [23, 41], but not yet specifically in redox oscillations. Our choice of modeling this system with positive twist is based on the results of our prior deterministic simulations. Measures of how fast mitochondrial oscillators respond to perturbations (the *λ* value in our simulations) in biochemical or cell culture assays are likewise rare. But in essence, simplifying the kinetic deterministic model into a stochastic amplitude-phase model allowed us, regardless of specific kinetic details and with the aforementioned assumptions, to quantitatively analyze the emergent properties that arise from a network of stochastic clocks with different intrinsic oscillator properties in response to coupling.

The presence of noise in cellular networks is a fundamental consequence of the stochastic nature of biochemical reactions. The ability to ensure proper transmission of information albeit such random fluctuations is a major issue in network behavior. In this paper, we have studied how different factors affect a simple model for redox circadian rhythms. We report and quantify both how noise and oscillator heterogeneity desynchronize a network of oscillators, and how inter-oscillator coupling and coupling to a TTFL helps in conferring resistance to noise, resulting in the generation of robust and high amplitude rhythms. This work highlights the importance of coupling in overcoming the intrinsic randomness of biochemical reactions and in contributing to robust circadian rhythmicity.

## V. Materials and Methods

### A network of stochastic amplitude-phase redox oscillators

Amplitude-phase oscillators are one of the most abstract yet intuitive class of models, with only two variables, amplitude and phase, that relate to oscillatory dynamics. The equations do not depend on specific molecular details and can describe phenomenologically the rhythmic behavior of a system [29, 37]. The stochastic differential equations of an individual Poincaré amplitude-phase redox oscillator *i*, in Cartesian coordinates, read

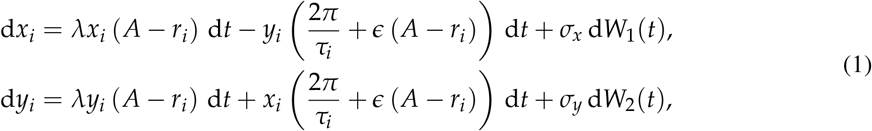

where 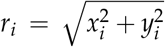; *A* represents the amplitude of the *D*_2_ oscillator (*A* = 1); *λ*, the amplitude relaxation rate (i.e., a measure of how fast a perturbation relaxes back to the limit cycle [29, 30, 52, 78, 81], *λ* = 0.05 h^-1^, Supplementary Figure S1B); *τ* the intrinsic period of the redox oscillator (*τ* = 24.23 h, Supplementary Figure S2C and also [19]) and *ϵ*, the oscillator’s explicit twist (*ϵ* = 0.05 h^-1^, Supplementary Figure S1A). Given the limited availability of quantitative data in mitochondrial redox oscillations, we explored the effects of different variances of Gaussian noise (Supplementary Figure S2A, B). In most of the simulations (unless stated), noise was introduced by adding a Wiener process d*W*(*t*) with *σ*_*x*_ = *σ*_*y*_ = 0.05 in the velocity field, thus modeling intrinsic noise. Estimation of the model parameters *λ, 20AC* and *τ* is described the following section.

To model how coupling affects an ensemble of *N* noisy *D*_2_ oscillators, we assumed that the individual oscillators *i* interact with each other through a mean-field *M*, defined as

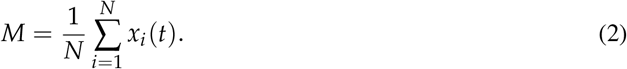

This kind of coupling has been used in previously published models of the circadian clock that study synchronization among oscillators [29, 53], and implies a relatively fast diffusion of coupling agents (H_2_O_2_ in the cytoplasm) compared to the ∼24 h circadian time scale. The mean-field *M* is then assumed to act directly on each *x*_*i*_ oscillator at a coupling strength *K*_*c*_ such that the network dynamics in the presence of coupling then read

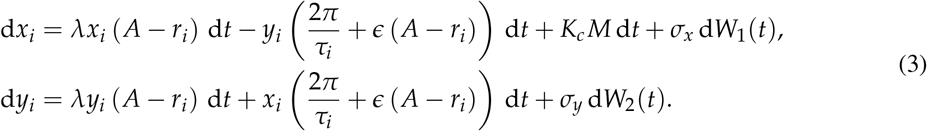

To quantify the re-organization of oscillator phases induced by increasing coupling strengths *K*_*c*_ we used the order parameter or synchronization index *R*_0_ [50, 82], defined as

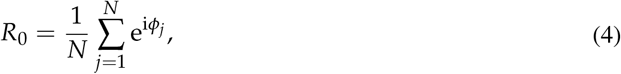

from the instantaneous phases and amplitudes of individual oscillators *j*.

### Estimation of amplitude-phase model parameters

The model parameters *τ, A* and *λ* were determined as follows. The free running period of *D*_2_ oscillations in the kinetic ordinary differential equation model was 24.23 h (Figure 1B, Supplementary Figure S2C). The amplitude *A* was set to 1 for convenience, following previous theoretical studies [29, 30, 52]. The amplitude relaxation rate *λ* was determined from exponential fits after simulating 100 perturbations on the deterministic *D*_2_ oscillations. More specifically, we introduced a pulse-like perturbation (mimicking, for example, an injection of H_2_O_2_ into a culture medium) at a random phase and let the oscillator relax back to the limit cycle. The relaxation rate was computed as the mean of the decay rates after fitting exponentially decaying curves to the maxima of 100 randomly perturbed time series, resulting in an average value of *λ* = 0.050 *±* 0.002 h^-1^ (see Supplementary Figure S1B for a representative fit).

The twist parameter *ϵ* was determined by analyzing the amplitude-period correlation of simulations in which the parameter that describes the leakage of *D*_1_ to the cytosol in the kinetic model was varied *±* 10% around its default value to mimic oscillator heterogeneity. This parameter was chosen over the others because it was found to be the parameter that controlled the oscillation period to the largest extent in our previous model [19]. We found that changes this parameter affected period and amplitude of *D*_2_ oscillations with both rhythmic parameters being positively correlated (Supplementary Figure S1A) and thus set the twist value to a representative value of *ϵ* = 0.05 h^-1^.

The variances of the Gaussian noise terms could not be estimated due to the lack of experimental data, but after exploring the effects that different noise variances have on the desynchronization of the average signal (Supplementary Figure S2A, B), we heuristically set it to *σ*_*x*_ = *σ*_*y*_ = 0.05.

### Coupling of redox rhythms to a Goodwin-like model of the TTFL

To model the effect of a TTFL input in an ensemble of weakly-coupled redox oscillators, we used the following Goodwin-like model [53]:

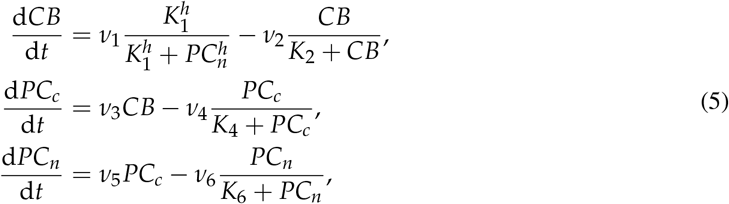

where *CB* represents a CLOCK:BMAL1 complex; *PC*_*c*_, the cytoplasmatic PER2:CRY1 complex after CLOCK:BMAL1-driven expression and translation; and *PC*_*n*_ represents the nuclear PER2:CRY1 complex that represses CLOCK:BMAL1 activity. Parameter values are taken from [53]: *ν*_1_ = 0.7 a.u. of conc./h; *K*_1_ = 1 a.u. of conc.; *h* = 4; *ν*_2_ = 0.35 a.u. of conc./h; *K*_2_ = 1 a.u. of conc.; *ν*_3_ = 0.7 h^-1^; *ν*_4_ = 0.35 a.u. of conc./h; *K*_4_ = 1 a.u. of conc.; *ν*_5_ = 0.7 h^-1^; *ν*_6_ = 0.35 a.u. of conc./h; *K*_6_ = 1 a.u. of conc.

Then, to simulate CLOCK:BMAL1-regulated generation of H_2_O_2_ [55–58], we added a *CB* term on the *y* variable of our weakly-coupled amplitude-phase redox ensemble as follows:

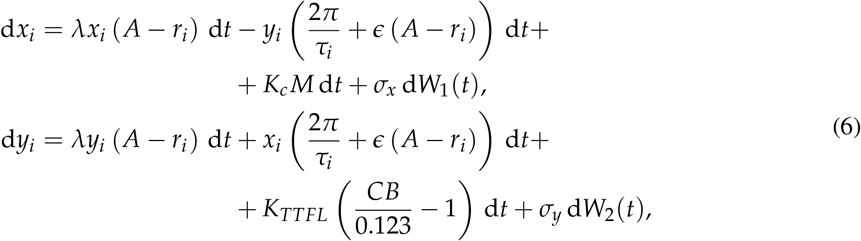

where *K*_*TTFL*_ represents the strength of the TTFL interaction; *M*, the mean-field as in equation 2; and *K*_*c*_, the strength of the inter-redox-oscillator coupling, which we set to 0.05 to simulate a weakly-coupled redox system. Since the *CB* solution obtained from equation 5 represents absolute levels of the CLOCK:BMAL1 complex, we normalized the solution by dividing it by its mean (0.123) and subtracting 1. This way, the time series of the redox amplitude-phase oscillator and of CLOCK:BMAL1 were both oscillating around 0.

### Numerical simulations

Results were obtained by solving the stochastic differential equations using the Euler-Maruyama method [38, 39] for a total integration time of 100 days at a Δ*t* = 0.01 h. At each time step, we introduced the white noise component by adding a random number in the differential equations of the *x*_*i*_ and *y*_*i*_ coordinates with standard deviations *σ*_*x*_ = *σ*_*y*_ = 0.05.

The Euler-Maruyama solver is available with the code in GitHub (https://github.com/olmom/coupled-oscillators-redox) and has been validated with ODE-based deterministic models.

## Abbreviations

ODE: ordinary differential equation
TTFL: transcription-translation feedback loop
Prx: peroxiredoxin
Srx: sulfiredoxin
*D*_1_: danger 1 (mitochondrial H_2_O_2_)
*D*_2_: danger 2 (cytosolic H_2_O_2_)
*A*: active peroxiredoxin
*I*: inactive peroxiredoxin
*R*: rescuer (mitochondrial sulfiredoxin).

## Acknowledgments

We thank Saskia Grabe and Gianmarco Ducci for fruitful discussions and technical help.

## Conflict of interest

The authors have no conflict of interest to declare.

## Data accessibility

The source code to generate the simulated data and reproduce all figures is available through GitHub (https://github.com/olmom/coupled-oscillators-redox).

## Funding

This study was supported by Deutsche Forschungsgemeinschaft (DFG, German Research Foundation) Project-ID 278001972 – TRR 186 to H.H., A.Kr. and M.dO.; SCH 3362/2-1 as well as SCH 3362/4-1 to C.S.

## Author contributions

Study design and conceptualization: M.dO. and H.H.; Methodology: M.dO. and H.H.; Investigation: M.dO. A.Ka. and H.H.; Resources and data curation: M.dO. and H.H.; Writing (original draft): M.dO. and H.H.; Writing (review & editing): M.dO., C.S., A.Kr. and H.H.; Funding acquisition: A.Kr. and H.H.

## Notes

https://github.com/olmom/coupled-oscillators-redox

### Competing Interest Statement

The authors have declared no competing interest.

