## Supplementary Figures S1-S5 for "Coupling allows robust redox circadian rhythms despite heterogeneity and noise"

### Supplementary Material

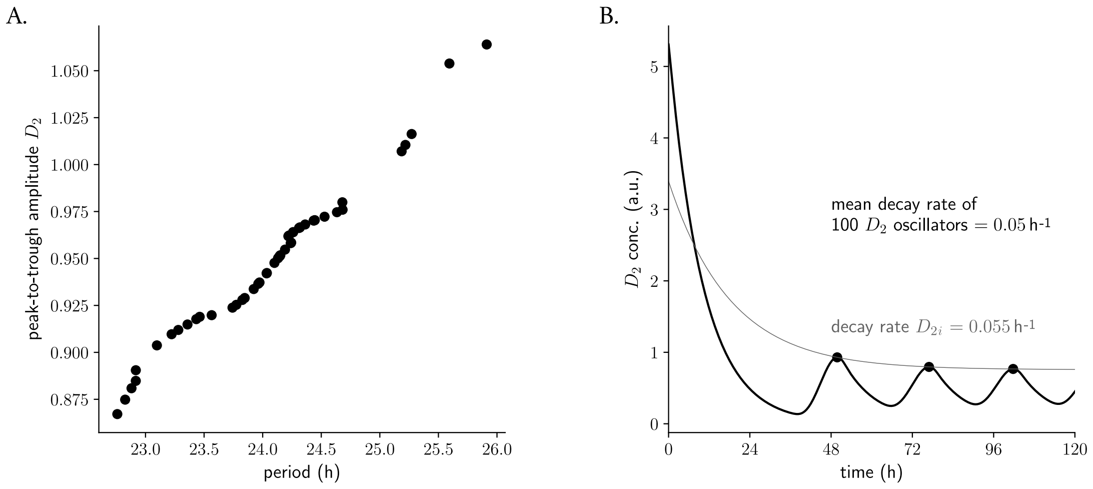


**Figure S1: Parametrization of the stochastic amplitude-phase $D_2$ oscillator. A.** Amplitude-period correlation (twist) in a network of heterogeneous $D_2$ oscillators. Heterogeneity was simulated by changing the parameter that represents the $D_1$ translocation to cytosol (where it is called $D_2$, see Figure 1A) in the kinetic model around $\pm 10\%$ of its default value $d = 0.2$ [19]. **B.** Representative decay upon a perturbation of $D_2$. To estimate the amplitude relaxation rate, perturbations were applied at different (random) phases of the $D_2$ time series and exponential decay functions were fitted to the maxima of the relaxation dynamics. The amplitude relaxation rate was computed as the average decay rate from 100 simulated perturbations.

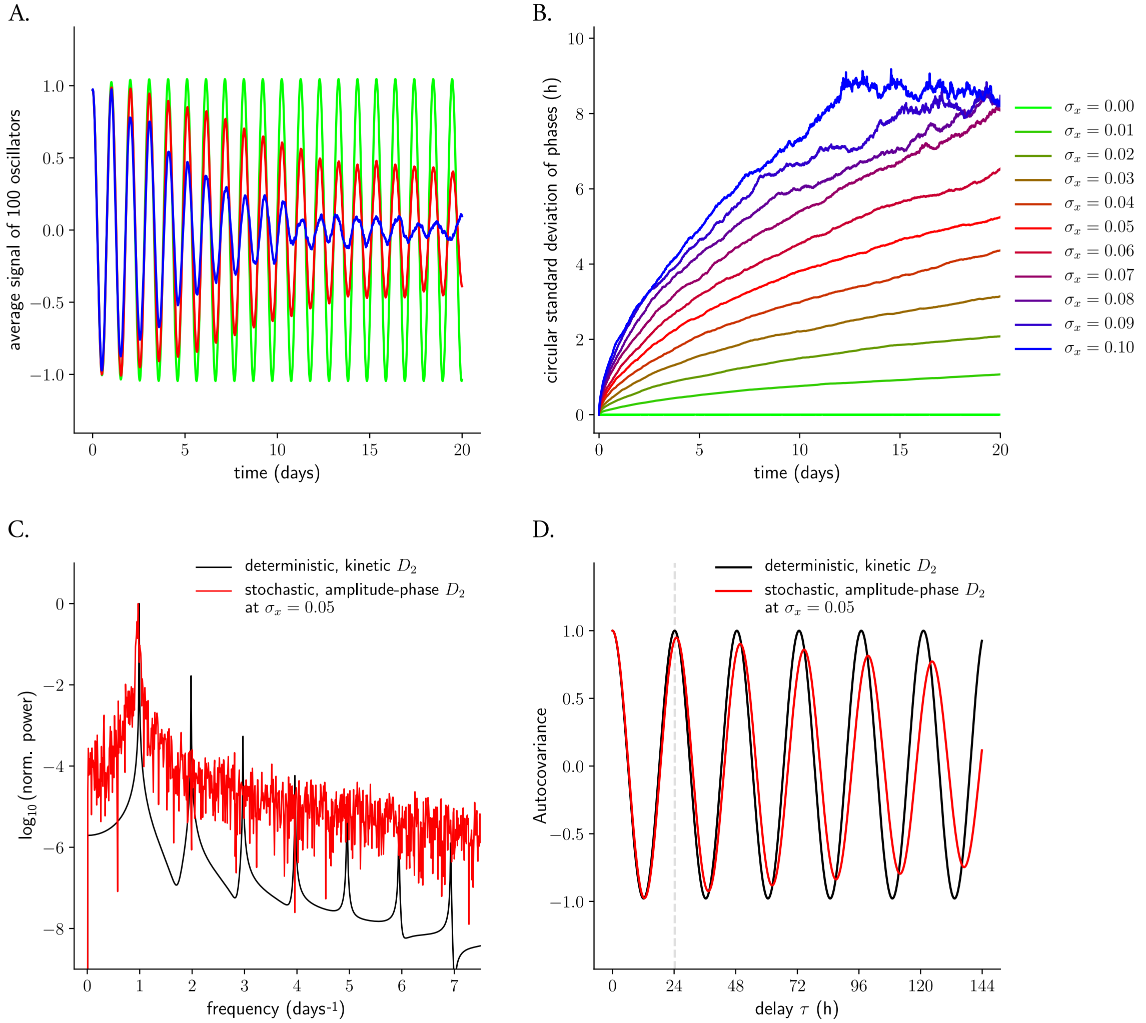


**Figure S2: Effect of noise on amplitude-phase $D_2$ oscillators. A.** Average signals from ensembles of 100 noisy amplitude-phase redox oscillators for different noise variances. $\sigma_x = 0.00$ in shown in green: the average signal of the network does not decay over time. $\sigma_x = 0.05$ and $\sigma_x = 0.10$ are shown in red and blue, respectively, and it is seen how the average signal decays over time. **B.** In networks of amplitude-phase oscillators, noise induces spreading of the phases of the individual oscillators: the higher the noise variance, the faster the phase-spreading. **C.** Spectral analysis of the $D_2$ time series from the kinetic ODE-based model (black) and from the stochastic amplitude-phase model at $\sigma_x = 0.05$ (red), to confirm that both models oscillate at the same frequency. The power was normalized to the maximum power. **D.** A representative example of the autocovariance of a deterministic, kinetic $D_2$ oscillator (black line, does not decay over time) and of a stochastic amplitude-phase oscillator at $\sigma_x = 0.05$ (red line, decays over time due to the noise component).

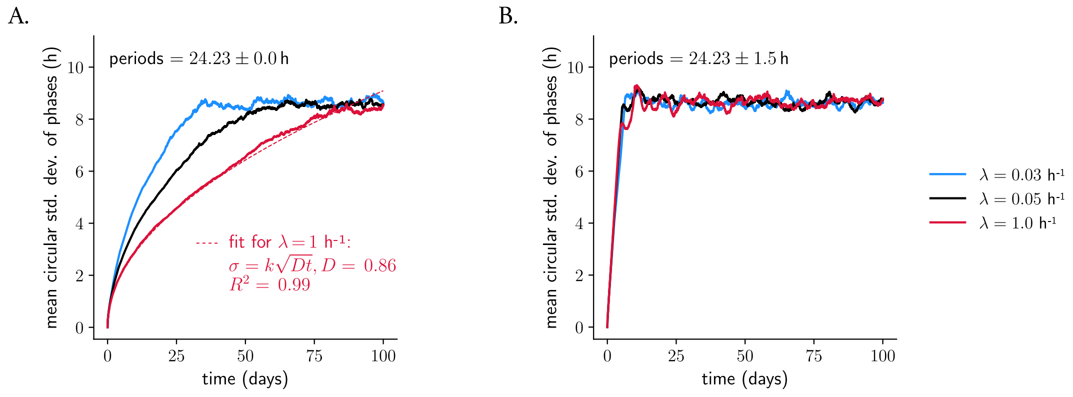


**Figure S3: Effect of amplitude relaxation rate on desynchronization dynamics in networks of identical vs. heterogeneous redox oscillators. A.** Effect of amplitude relaxation rates on desynchronization dynamics when all oscillators are identical: weaker oscillators (lower $\lambda$ values, blue and black curves) result in faster phase dispersion dynamics over time, that no longer fit to Fick's expected square root law. The phase dispersion grows with the square root of time only for rigid oscillators ($\lambda = 1\,\text{h}^{-1}$, red curve, with a diffusion coefficient $D = 0.86$ –red dashed line). **B.** Amplitude relaxation rate effects are masked with oscillator period heterogeneity. All oscillators dephase with roughly the same dynamics, independent of amplitude relaxation rate values, when oscillator periods are taken from a normal distribution at $24.23 \pm 1.5$ h. The average standard deviation from 50 different realizations of network simulations (each composed of 100 oscillators) is shown in both panels.

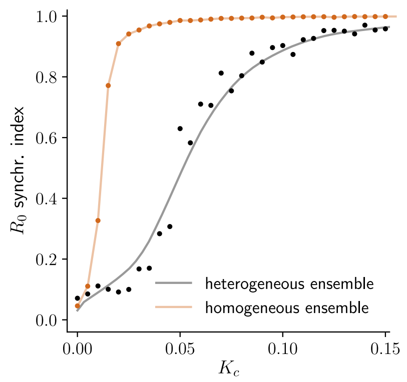


**Figure S4: Effect of oscillator heterogeneity on coupling-induced synchronization of an ensemble of stochastic amplitude-phase redox oscillators.** Phase synchrony is achieved for lower values of coupling strengths when the network is composed of identical oscillators (no period differences, orange curve). A heterogeneous network (with period differences, black curve) needs higher $K_c$ values for phase synchronization. Note though, that although $R_0 = 1$ is not reached in the case of a heterogenous oscillator ensemble for $K_c = 0.10$, they can be synchronized with respect to periods (see Figure 3A, lower panel). Points indicate the mean $R_0$ from 5 different realizations of network simulations (each realization is composed of a network of 100 oscillators); the solid black line represents the average of 100 Lowess models that were fitted to 50% of the points (chosen randomly in each Lowess iteration).

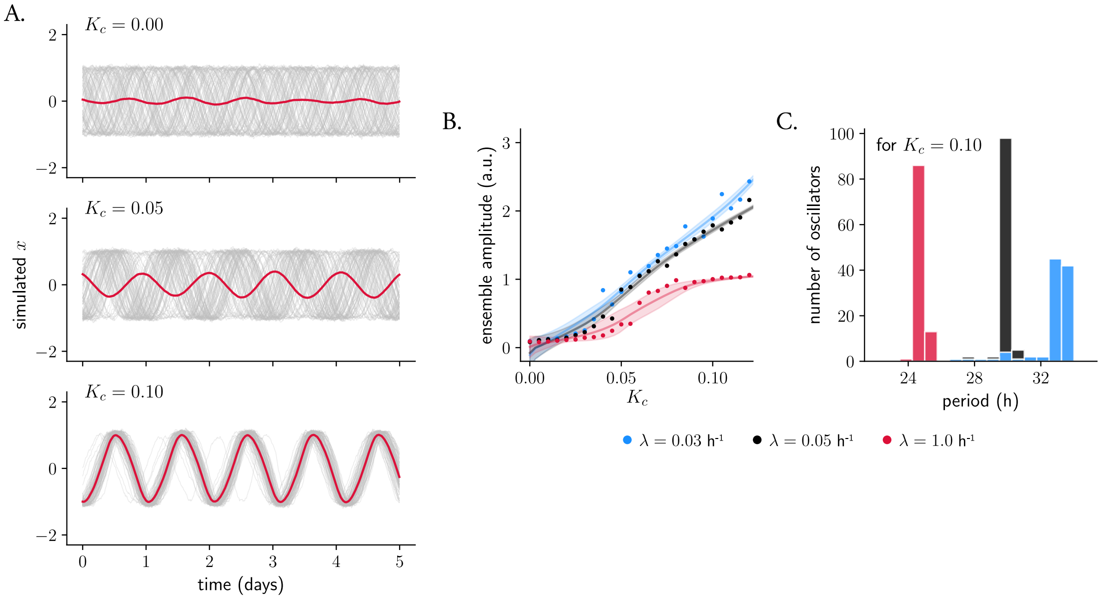


**Figure S5: Effect of amplitude relaxation rate in coupling-induced amplitude and period expansions due to twist.** **A.** Coupling induces temporal order in a network of 100 rigid stochastic redox amplitude-phase oscillators ($\lambda = 1\,\text{h}^{-1}$). Individual oscillators are shown in grey; the mean-field is shown in red. **B.** Effect of amplitude relaxation rate in coupling-induced amplitude expansions of the mean-field. Weaker oscillators (lower $\lambda$ values, blue and black curves, $\lambda = 0.03\,\text{h}^{-1}$ and $0.05\,\text{h}^{-1}$, respectively) respond with larger amplitude expansions upon coupling than more rigid oscillators (red curve, $\lambda = 1\,\text{h}^{-1}$). Points indicate the mean ensemble amplitude of 5 different realizations of network simulations; the solid line represents the average of 100 Lowess models that were fitted to 50% of the points (chosen randomly in each iteration); the lighter shaded areas indicate the confidence interval for the 100 Lowess models. **C.** Effect of amplitude relaxation rate in coupling-induced period expansions of the mean-field. Weaker oscillators (lower $\lambda$ values, blue and black bars, $\lambda = 0.03$ and $0.05\,\text{h}^{-1}$, respectively) are less robust to period expansions for a representative $K_c = 0.10$ than more rigid oscillators (red bars, $\lambda = 1\,\text{h}^{-1}$). These expansions can be explained with twist: rigid oscillators are more robust to twist-induced effects.
